# Integrated meditation and exercise therapy: A randomized controlled trial of a combined non-pharmacological intervention reduces disability and pain in patients with chronic low back pain

**DOI:** 10.1101/652735

**Authors:** Anna M. Polaski, Amy L. Phelps, Thomas J. Smith, Eric R. Helm, Natalia E. Morone, Kimberly A. Szucs, Matthew C. Kostek, Benedict J. Kolber

## Abstract

Integrative and complementary non-pharmacological treatments have proven efficacious in treating both the physiological and psychological symptoms of chronic pain conditions but the potential of many combined therapies is unknown. This study examined the effects of a combined intervention of mindfulness meditation followed by aerobic walking exercise in chronic low back pain (cLBP) patients. We hypothesized that meditation before exercise would reduce disability and pain by increasing mindfulness prior to physical activity. Thirty-eight adults completed either meditation and exercise treatment (MedExT) (n=18) or an audiobook control condition (n=20). Over a 4-week period, participants in the MedExT group performed 12-17 minutes of guided meditation followed by 30 minutes of moderate intensity walking exercise 5 days per week. Measures of disability, pain, mindfulness and anxiety were taken at baseline and post-intervention. Ratings of pain were also assessed by participant self-report, before and after each intervention session. Following MedExT, participants showed significant improvement in our primary outcome of disability compared to the control group (p<0.05). From pre to post-intervention, MedExT also increased mindfulness (p<0.05), but had no significant effect on quantitative sensory testing on the low back. Mean ratings of low back pain intensity and unpleasantness significantly improved with MedExT from before the study to during participation, respectively (intensity p<0.05; unpleasantness p<0.05). Overall, four weeks of MedExT produced substantive changes in disability, mindfulness and measures of pain intensity and unpleasantness.

## INTRODUCTION

Low back pain is the most common reported type of pain [16] and the second leading cause of physician visits and disability among U.S. adults [19]. Globally, 25% of adults report having low back pain over any one month [27]. Often due to non-specific causes and complicated by comorbid symptoms [19], low back pain remains difficult to treat. Current treatments include NSAIDS, muscle relaxants, opioids, psychological therapy, physical therapy chiropractic manipulation, injections and surgery [1, 19]. Chronic low back pain is further made complex by the potential for comorbid anxiety disorders [34]. In particular, musculoskeletal pain is often associated with fear avoidance anxiety behavior and kinesiophobia [35]. This kinesiophobia or fear of movement can further exacerbate pain and subsequent disability. Kinesiophobia may also reduce the potential benefit of physical treatments in patients by increasing state anxiety before and during therapy. In treating chronic pain, a major gap exists in not only treating the physiological condition, but also addressing the interplay with psychological etiologies.

Due to the risk of adverse side effects, addiction and misuse, many pharmacological approaches to treating low back pain, including opioid therapeutics, have not been found to be superior to complementary treatment methods [41]. There has been a significant push in the last 20 years to identify and understand complementary and integrative therapies to supplement pharmacology. Nonpharmacological therapies include aerobic exercise, tai chi, yoga, mindfulness-based stress reduction (MBSR), progressive relaxation, electromyography biofeedback, operant therapy, cognitive behavioral therapy, multidisciplinary rehabilitation, acupuncture, spinal manipulation and massage with many of these showing significant positive effects [13]. There has been considerable interest in programs that combine elements of multiple therapies to treat chronic pain. One of most well-established integrative programs that involves elements of stress reduction, exercise, and meditation is the 8-week MBSR system, which has been found to improve pain, depression and quality of life [26]. However, this program requires extensive training and may not be easily accessible to some persons with cLBP. In the current study, we examined a more feasible pilot program of introductory mindfulness meditation that novice meditators could easily put into practice prior to aerobic walking exercise. Both meditation and exercise have been independently investigated in the context of back pain therapy.

Exercise interventions have proven to have beneficial outcomes on pain severity, physical disability, psychological function and health-related quality of life across various chronic pain conditions [23]. Mechanistically, aerobic exercise at a level of at least 70% of the maximum aerobic capacity generates the production of endorphins and elicits other pain inhibitory mechanisms driven by the central nervous system [6, 37]. In addition, aerobic exercise has been shown to reduce fatigue and improve peak oxygen uptake, and physical fitness [17, 25].

Similar to exercise, studies incorporating mindfulness meditation have largely shown to improve pain and depression symptoms, quality of life, well-being and increase mobility and functioning [26, 36]. Mechanistically, meditation with mindfulness has been associated with decreased levels of cortisol [28], increased signaling connections in the brain [49], improved pain processing and emotional control [31], and altered amygdalar response to emotional stimuli [15]. As these therapies (exercise and meditation) independently improve disability and pain, the <u>Med</u>itation and <u>Ex</u>ercise to <u>T</u>reat chronic low back trial (MedExT) tested the effects of a 4-week intervention of a guided mindfulness meditation program combined with moderate intensity walking exercise performed 5 days per week in chronic low back pain (cLBP) patients. We hypothesized that this intervention would improve disability (primary outcome), pain, anxiety and increase mindfulness compared to control participants. This specific therapy combination has not been previously examined in chronic pain patients.

## METHODS

### Participants

Participants were 52 adults (age 18-60) with chronic low back pain (>6 months) with no evidence of neuropathic pain, radicular pain (i.e. sciatica), or referred somatic pain. Participants were recruited using in-clinic recruitment to the University of Pittsburgh Department of Physical Medicine and Rehabilitation Research Registry (PMR3) and the University of Pittsburgh Clinical and Translational Science Institute (CTSI) patient registry, Pitt+Me. Initial pre-screening was completed during recruitment to the Pitt+Me database with phone follow-up by A.M.P. Full inclusion criteria included 1) a BMI within the normal to overweight range (18.5-29.9), 2) a resting heart rate between 60 and 100 bpm, 3) resting blood pressure less than or equal to 140/90mmHg, and 4) the ability to independently ambulate community distances without external support (i.e. walker, cane). Exclusion criteria included 1) cardiovascular or respiratory disease, 2) neurological disease, unrelated to low back pain, 3) diabetes mellitus, Types 1 and 2, 4) diagnosis of chronic pain condition unrelated to low back pain, 5) acute pain, 6) regular participation in high intensity athletic/sporting activities, 7) sedentary lifestyle, 8) currently pregnant individuals, 9) current cigarette smokers, 10) individuals with on-going litigation associated with back pain, 11) regular participation in meditation techniques or training in mindfulness-based stress reduction.

### Study Design

This study was designed as a randomized single-blinded (for QST testing) controlled trial with repeated measures testing the effect of a combined treatment of mindfulness meditation and aerobic walking exercise (MedExT) compared to a control intervention. The trial was randomized between the two groups using a random sequence generator. M.K. was responsible for generating random allocation sequence and A.M.P. was responsible for enrolling and assigning participants to interventions. QST outcome assessor (B.J.K.) remained blinded to treatment assignment. A power analysis indicated that a minimum of 21 subjects/group to be sufficient to detect statistical differences in our primary dependent variable, disability measured with the Roland Morris Disability Questionnaire (RMDQ) (alpha=0.05, effect size=0.8, power=0.80) using the G-power calculator based on previously published work using MBSR and low-back pain. All procedures were approved by the Duquesne University Institutional Review Board (Protocol #2017-05-12) and written consent was obtained from each participant prior to testing. All methods were performed in accordance with the relevant international and local guidelines and regulations for human research. This study is registered with ClinialTrials.gov under ID: NCT03324659 (10/30/2017). Participants were compensated for participation.

### Procedures

In-clinic sessions were conducted at Duquesne University’s Exercise Physiology Laboratory over the course of the 4-week intervention period between January 2018 to April 2019. For participants meeting phone screening criteria, informed consent was obtained and participants were enrolled in the study. An initial clinical screening exam was performed by three clinicians (E.H. or two trained Physician Assistants). During this screening (∼15 minutes), patients were evaluated for strength, lumbar range of motion, reflexes and sensation in relation to their low back pain. This screening was done to verify back pain inclusion (e.g. exclude radicular patients) and to determine safety of participation in the exercise portion of the intervention. Of 55 patients recruited, no patients were excluded during this screening. Following clinical screening patients were scheduled to start the actual intervention. The average time between consent and start of trial was 26 days. At the start of the full trial (after clinical screen), participants came in for an intake session during which they completed a battery of questionnaires (see *Instruments* below) and were oriented to the general study protocol. The intake session consisted of a sequence of quantitative sensory tests and baseline assessments of pain (see *QST* section below). B.J.K. performed all QST blinded to the treatment group of the participants and remained blinded to treatment until after the final pain assessments were completed. Participants were blinded to treatment assignments for baseline intake testing. Following baseline testing, treatment assignments were disclosed to the participants.

Within one week of performing baseline pain assessments (average time between baseline and first intervention session = 5 days), participants completed their first in-clinic intervention session. At the start of this session, patients received approximately 35-45 minutes of meditation or stress training by a clinical psychologist (T.S.). These sessions discussed either the potential of and use of mindfulness and meditation (MedExT group) or general stress management and wellbeing for chronic pain (control group). Sessions were standardized by using a script developed by T.S. (see document, **Supplemental Digital Content 1**). Following this session, subjects completed their first intervention session, either combined meditation and exercise (MedExT) or the control condition. Participants had the option to complete intervention sessions at-home or in-clinic. Interventions were performed 5 days per week for 4 weeks. In-clinic intervention sessions were typically attended once per week. During these sessions two experimenters were present and did a check-in with the participants to ensure that they were not experiencing any difficulty completing the assigned intervention. 48 hours after the end of the 4-week period, participants attended the exit session, where they again completed surveys and underwent QST.

### Meditation and Exercise Protocol

For subjects in the MedExT experimental group, guided meditation recordings with a focus on mindfulness by meditation teacher and psychologist Dr. Tara Brach were used [9, 22]. A selection of five different recordings were utilized; each recording was listened to one time per week and lasted between 12-17 minutes (see **Supplemental Digital Content 2** for URLs to recordings). Recordings were selected by T.S. along with clinical psychologist Ian C. Edwards for their focus on mindfulness and overall length. Participants were given an mp3 player (SanDisk) loaded with each of the five meditation recordings to borrow. During the weekly in-clinic session, subjects practiced the meditation portion of the intervention session in our interdisciplinary meditation room which was a quiet space with low lighting and comfortable seating options. For at-home intervention sessions, subjects were encouraged to perform meditation in a quiet comfortable setting.

Immediately following meditation, participants performed 30 minutes of moderate intensity walking exercise on a treadmill. Prior to the first exercise session, resting heart rate and age was used to calculate a heart rate that corresponded to 50% heart rate reserve (HRR) for each participant [44]. We used the 50% HRR estimate as the target heart rate for moderately intense exercise with a range of 40-60% HRR calculated for each participant. Heart rate monitors (Polar H1) were worn for each in-clinic exercise session to monitor exertion levels. During the first in-clinic exercise session, trial coordinator A.M.P would manipulate the speed and grade of the treadmill in order to achieve the calculated heart rate for an individual participant. Average grade was 2.4% and speed range was 2.2-3.8 mph. Once reached, this speed and grade combination was used as the walking prescription for subsequent exercise sessions for that particular participant. Prior to and following exercise, each MedExT experimental intervention participant rated their perceived exertion levels using the Borg RPE scale [8]. Each exercise session began with a 2-minute warm-up at 2.5 mph and concluded with a 2-minute cool-down (total time 30 minutes on treadmill).

### Control Protocol

Participants in the control group listened to an audiobook for 12-17 minutes followed by a 30-minute rest period 5 times per week for 4 weeks. Each session was time-matched to the experimental intervention group. Subjects were given an mp3 player with 20 (one for each day) recordings of The *Natural History and Antiquities of Selborne* [54], which has been previously used and validated as a neutral comparison for guided relaxation interventions [14, 51]. During the resting period, participants were free to read, watch television, listen to music or other activity that was less than moderate physical effort and not stressful.

### Survey Instruments and Administration

All surveys were administered using Qualtrics XM Research Core software [45] either via a tablet for in-clinic sessions or via email for at-home sessions. All subjects completed the following questionnaires at baseline and exit: the Roland Morris Disability Questionnaire (RMDQ) [46], the State-Trait Anxiety Inventory (STAI form Y) [48], and the Fear-Avoidance Beliefs Questionnaire (FABQ) [52]. The AHA/ACSM Pre-participation Screening Form [3] and the International Physical Activity Questionnaire (IPAQ-short) [7] were also completed at baseline to assess eligibility for enrollment. The Freiburg Mindfulness Inventory (FMI) [53] was administered prior to the mindfulness training session and again at the exit session for MedExT experimental intervention subjects.

Pain was assessed using quantitative sensory testing methods (described below), as well as self-report measures of pain using a visual analog scale (VAS) consisting of a 10cm line with the numbers 0 and 10 at either end for intensity and unpleasantness ratings. On each day of the assigned intervention, participants received reminder emails with a URL link to the daily VAS survey, on which subjects would rate pre and post-intervention VAS pain intensity and unpleasantness. This survey was able to capture time stamps of survey progress, allowing for monitoring of protocol compliance.

Throughout the 4-week trial period, participants in both groups wore ActiGraph GT9X Link devices in order to monitor physical activity (steps per day). During the exit session, participants were also given an exit survey that was used to identify likelihood of continued adherence (for MedExT group) and any barriers to this intervention. This survey was qualitatively analyzed.

### Quantitative/Qualitative Sensory Testing (QST)

Quantitative sensory testing was done on the bare skin of the participant’s low back and forearms at specific testing sites. These assays assessed each participants’ cutaneous mechanical sensitivity (threshold for mechanical detection to 0.008g, 0.02g, 0.04g, 0.07g, 0.16g, 0.4g, 0.6g and 1.0g Touch Test filaments in 3 of 5 trials for filament), cutaneous mechanical pain (threshold for mechanical detection up to 300g Touch Test filaments), constant heat pain (45°C 3cm x 5cm heating block applied for 3 seconds followed by 10cm Visual Analog Scale (VAS) for intensity and unpleasantness of pain), pressure pain threshold (1cm round probe applied at constant ramping pressure until participant defined cutoff in kg at “pain threshold”; Wagner Instruments, Greenwich, CT, USA) and constant pressure pain sensitivity (2 second pressure stimulus at participant defined threshold followed by VAS for intensity of pain and unpleasantness of pain) as previously described [33]. 10cm VAS scales were numbered at 0 and 10. Score was measured to the nearest mm. Intensity scale ranged from 0=“*No pain”* to 10=“*The worst pain imaginable”*. Unpleasantness scale ranged from 0=“*Not unpleasant*” to 10=“*Most unpleasant sensation imaginable*”. Testing was performed at baseline and post-intervention to measure the overall change in sensitivity across the entire study.

### Statistical Analysis

Prior to analysis, an *a priori* statistical plan was developed (ClinialTrials.gov under ID: NCT03324659). Descriptive statistics were calculated using the IBM SPSS Version 25 and graphed with either SPSS or GraphPad Prism (Version 6.0). Normality of the data was assessed. Nonparametric inferential statistics were used for data that were not normally distributed. For analysis of primary and secondary outcomes, we were interested in looking at mean change, however the raw data values can be found as a table (see **Supplementary Digital Content 3**). *A priori*, we determined that participants had to complete 80% of the weekly sessions (≥ 4 of 5 sessions per week) for inclusion in data analysis.

#### Primary Outcome

The primary outcome was defined as the comparison of post-intervention RMDQ scores between the MedExT and the control groups. A two-sample t-test was used to identify a significant difference between groups using p<0.05. This questionnaire was chosen as the primary outcome measure because it was recently used in both a pilot study and large-scale assessment of MBSR in chronic low back pain patients [38, 39].

#### Secondary Outcomes

Five groups of secondary outcomes were measured and analyzed. P values were adjusted for each group of analyses based on the number of tests in that group. The first analysis tested whether the MedExT group would significantly increase mean scores on the Freiburg Mindfulness Inventory as determined by a two-sample t-test (p<0.05). The second explored whether the MedExT treatment would significantly influence a mean change in responses on three psychological inventories administered: 1) the Fear-Avoidance Beliefs Questionnaire, 2) the STAI state anxiety inventory and 3) the STAI state trait anxiety inventory. These were analyzed using two-sample t-tests where p<0.02 was considered significant. The third group of secondary outcomes measured mean response changes in the series of 14 QST taken at baseline and at the completion of the 4-week intervention period on the low back and non-dominant forearm. Analyses were grouped separately for tested body site. Significant mean pre/post differences between groups were identified using two-sample t-tests. Mann-Whitney-Rank-sum tests were used for mechanical sensitivity and mechanical pain at each site. Given the number of statistical tests (n=7 per body site) required for the QST secondary outcome measurements, a corrected p<0.005 was utilized for each body site to determine statistical significance.

Fourth, we assessed current back pain using VAS during each day of the trial. These VAS measurements were repeatedly made throughout the study taken at baseline and on each intervention day, pre and post-session. These measures were the VAS pain intensity score and the VAS pain unpleasantness score. In our original stats plan, a repeated measures MANOVA was to be utilized to determine if the vector of timed responses was significantly different between the two study groups. Due to missing data from some days, the MANOVA statistical plan was modified to using a mixed error-component model during analysis of data. JMP was used to perform this analysis. Participants were instructed to evaluate their on-going back pain at the time of the measurement. We looked at this in two ways: (1) the overall effect of the 4-week intervention on intensity and unpleasantness ratings (each day’s pre-intervention measurement minus baseline) and (2) the acute effect of each day’s session on VAS ratings (post-intervention VAS minus pre-intervention VAS).

Fifth, a final secondary outcome assessed “the average” pain that a participant experienced using intensity and unpleasantness VAS scales. During the exit session, participants evaluated VAS ratings of average low back pain intensity and unpleasantness that they remembered experiencing before the start of the study and after the 4 weeks of the intervention period. Significant mean response differences between groups were identified using two-sample t-tests. To correct for the number of comparisons, p<0.025 was considered a significant difference between groups.

#### Demographic Variables

The following demographic variables were collected and compared between groups to further check against potential bias: age, sex, handedness, body mass index (BMI), baseline heart rate (HR), baseline blood pressure (BP), baseline IPAQ-short, and mean number of steps taken per day over the 4-week intervention period. This was done using two-sample t-tests. Difference in the proportion of sex and handedness was tested using the Fisher’s Exact test where p<0.05 was considered significant. All other continuous variables were tested using two-sample t-tests for significant differences between the two study groups (p<0.05).

## RESULTS

### Participant Characteristics

Fifty-two adult volunteers with chronic low back pain were enrolled in this trial and thirty-eight participated in its entirety.14 participants dropped out of the study after enrollment. This included 10 due to scheduling conflicts, 3 due to newly discovered ineligibility (e.g. neurological disorder), and 1 due to inability to complete minimum required 80% sessions per week. See **Figure 1** for flow-chart diagram. Recruitment of participants began in January 2018 and ended February 2019. Demographic characteristics of subjects are presented in **Table 1**. Two-sample t-tests revealed no significant group differences for any of the demographic variables. A Fisher’s exact test found no significant relationships comparing treatment group to sex and also to handedness (p>0.05). Using ActiGraph watch data, we compared the average number of steps taken per day for participants in both groups. After subtracting steps taken by the MedExT group during their 30-minute exercise session, we found no statistically significant difference between the groups (p>0.05).

**Table 1.**
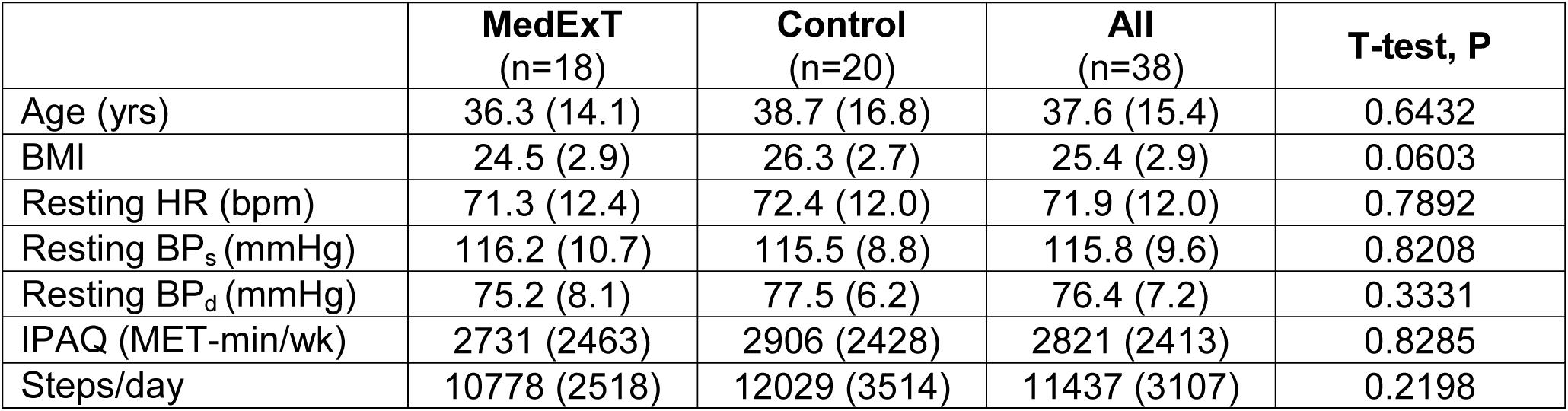
Participant characteristics. Data are mean (SD). Abbreviations: BMI, body mass index; BP_s_, systolic blood pressure; BP_d_, diastolic blood pressure; IPAQ, International Physical Activity Questionnaire.

|  | **MedExT**<br>(n=18) | **Control**<br>(n=20) | **All**<br>(n=38) | **T-test, P** |
| --- | --- | --- | --- | --- |
| Age (yrs) | 36.3 (14.1) | 38.7 (16.8) | 37.6 (15.4) | 0.6432 |
| BMI | 24.5 (2.9) | 26.3 (2.7) | 25.4 (2.9) | 0.0603 |
| Resting HR (bpm) | 71.3 (12.4) | 72.4 (12.0) | 71.9 (12.0) | 0.7892 |
| Resting BP<sub>s</sub> (mmHg) | 116.2 (10.7) | 115.5 (8.8) | 115.8 (9.6) | 0.8208 |
| Resting BP<sub>d</sub> (mmHg) | 75.2 (8.1) | 77.5 (6.2) | 76.4 (7.2) | 0.3331 |
| IPAQ (MET-min/wk) | 2731 (2463) | 2906 (2428) | 2821 (2413) | 0.8285 |
| Steps/day | 10778 (2518) | 12029 (3514) | 11437 (3107) | 0.2198 |

**Figure 1.**
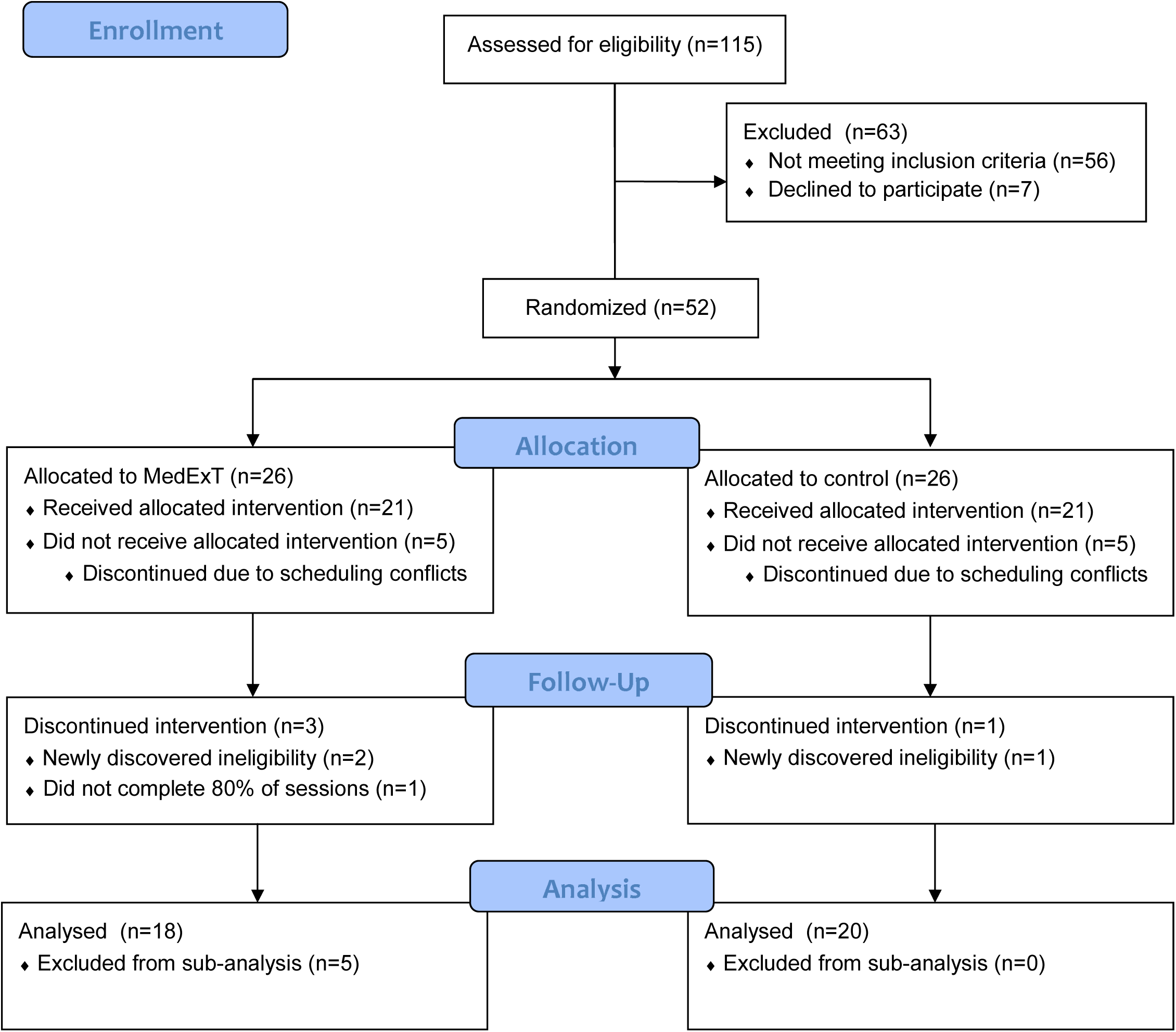
Consort Flow Diagram.

### Primary outcome: Intervention effects on Disability

Our primary outcome was the effect of treatment on post-intervention scores of disability as measured by the RMDQ. A two-sample t-test indicated a significant improvement in disability scores for the MedExT group compared to control (p=0.0123) (**Fig. 2**).

**Figure 2.**
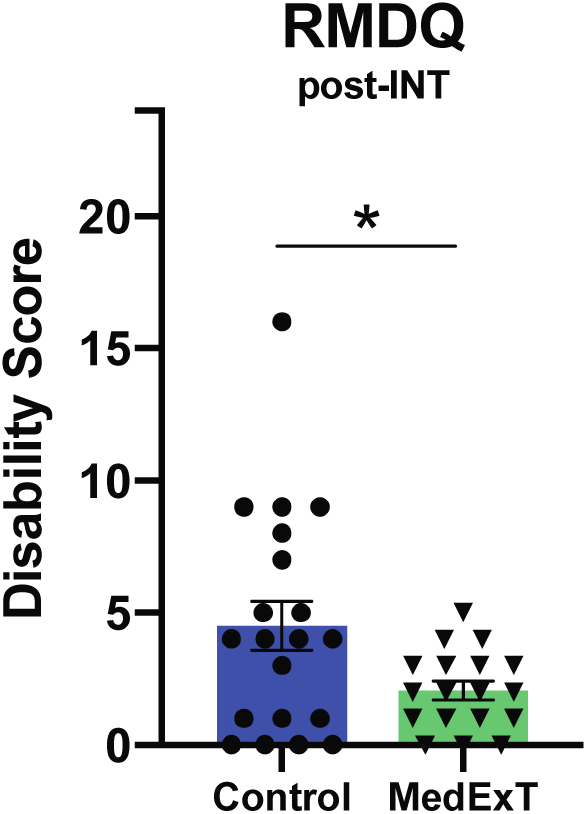
Post-intervention effect of MedExT vs. control treatment on primary outcome: disability as measured by the Roland Morris Disability Questionnaire (RMDQ). MedExT participants show statistically significantly lower disability levels compared to control participants. Data shown as mean +/− SEM. RMDQ min score=0, max score=24. *p<0.05.

### Secondary outcomes: Mindfulness, Fear Avoidance, Anxiety, and Pain

The FMI was administered to determine if there were any changes in mindfulness that developed during the trial. A two-sample t-test revealed a significant increase in mindfulness for the MedExT group from pre to post-intervention (p=0.0141) (**Fig. 3A**). For the psychological inventories, we tested whether the MedExT treatment would influence a mean change in response from pre to post-intervention. Two-sample t-tests showed no significant differences between pre and post measures for the MedExT group for the FABQ (p>0.02), STAI state anxiety (p>0.02), or STAI trait anxiety (p>0.02) (**Fig. 3B-D**).

**Figure 3.**
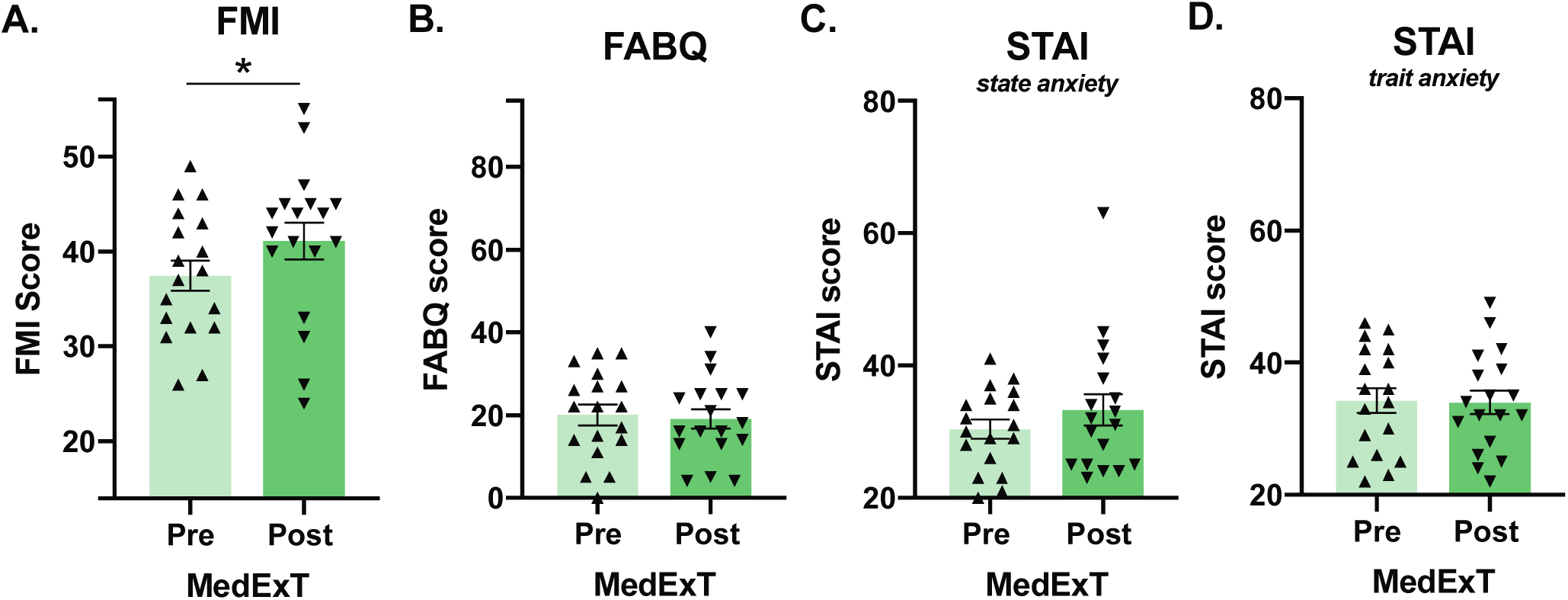
Pre to post-intervention effect of MedExT on secondary outcomes: (A) MedExT participants demonstrated statistically significant increases in mindfulness (FMI) (*p<0.05) compared to baseline values. No significant changes were observed for (B) fear avoidance behavior (FABQ) and (C) state and (D) trait anxiety (STAI). FMI score range=14-56, FABQ=0-96, STAI=20-80. Data shown as mean +/− SEM. For psychological inventories (FABQ, STAI), p>0.02.

For quantitative measures of pain (QST), we analyzed mean response changes from pre to post-intervention on the participant’s low back and non-dominant forearm to determine any significant differences between groups. Body site specific data for each test are shown in **Table 2**. For the low back and forearm, two-sample t-tests found no significant effects of treatment for constant heat pain intensity, constant heat pain unpleasantness, pressure pain threshold, constant pressure pain intensity or constant pressure pain unpleasantness (p>0.005). Additionally, Mann-Whitney tests showed no significant differences between treatment for mechanical sensitivity or mechanical pain for either the low back or forearms (p>0.005).

**Table 2.**
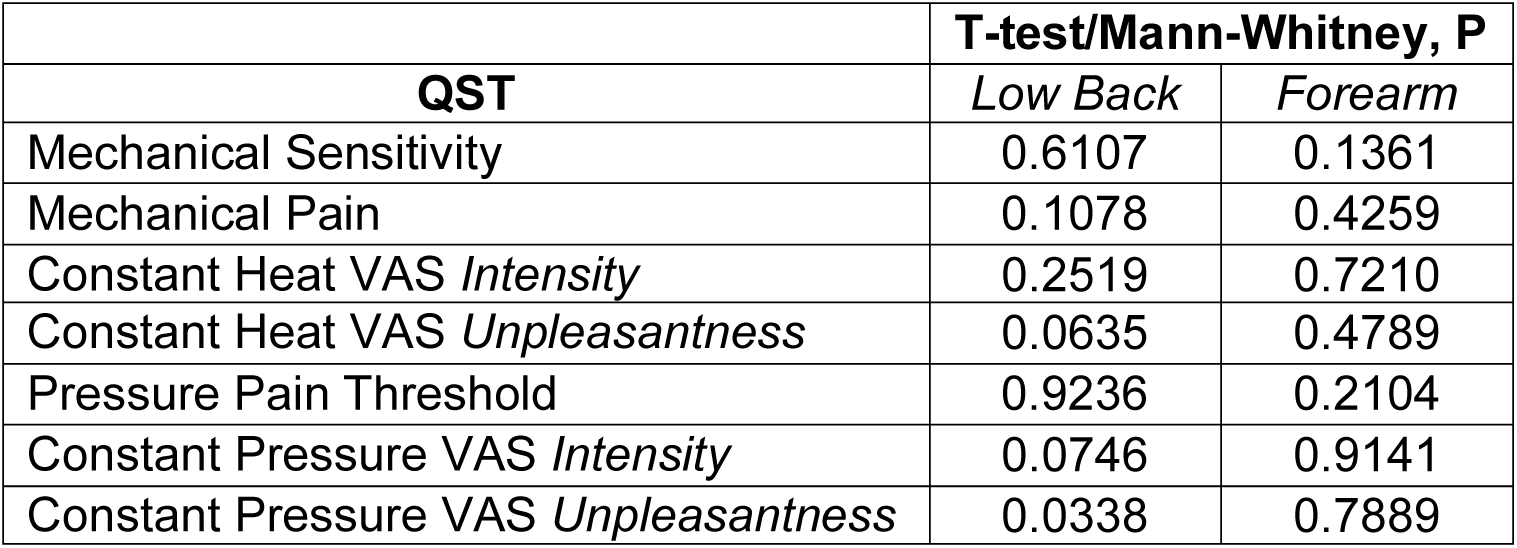
Intervention effects on QST pain measures for MedExT and control groups. For all QST, p>0.005.

For VAS repeated measures of on-going back pain, we found analgesic effects of the intervention that appear to accumulate over time (**Fig. 4**). A mixed-effects model revealed a significant effect of time (p=0.0008) and time x treatment (p=0.0012) for intensity ratings on each day before undergoing the intervention session (**Fig. 4A**). For unpleasantness ratings pre-intervention, a mixed-effects model showed significant effects of treatment (p=0.0330), time (p=0.0022) and time x treatment (p<0.0001) (**Fig. 4B**). Analysis of acute day to day effects of intervention indicated no significant effects for intensity (**Fig. 4C**), but a significant effect of treatment (p=0.0049) for unpleasantness post – pre measures (**Fig. 4D**). That is, the intensity VAS measured immediately after the ∼45-minute session was not significantly different from the VAS measured immediately before that day’s session. The lack of an effect here illustrates the potential cumulative effect of the intervention on pain rather than an acute exercise-induced hypoalgesia effect.

**Figure 4.**
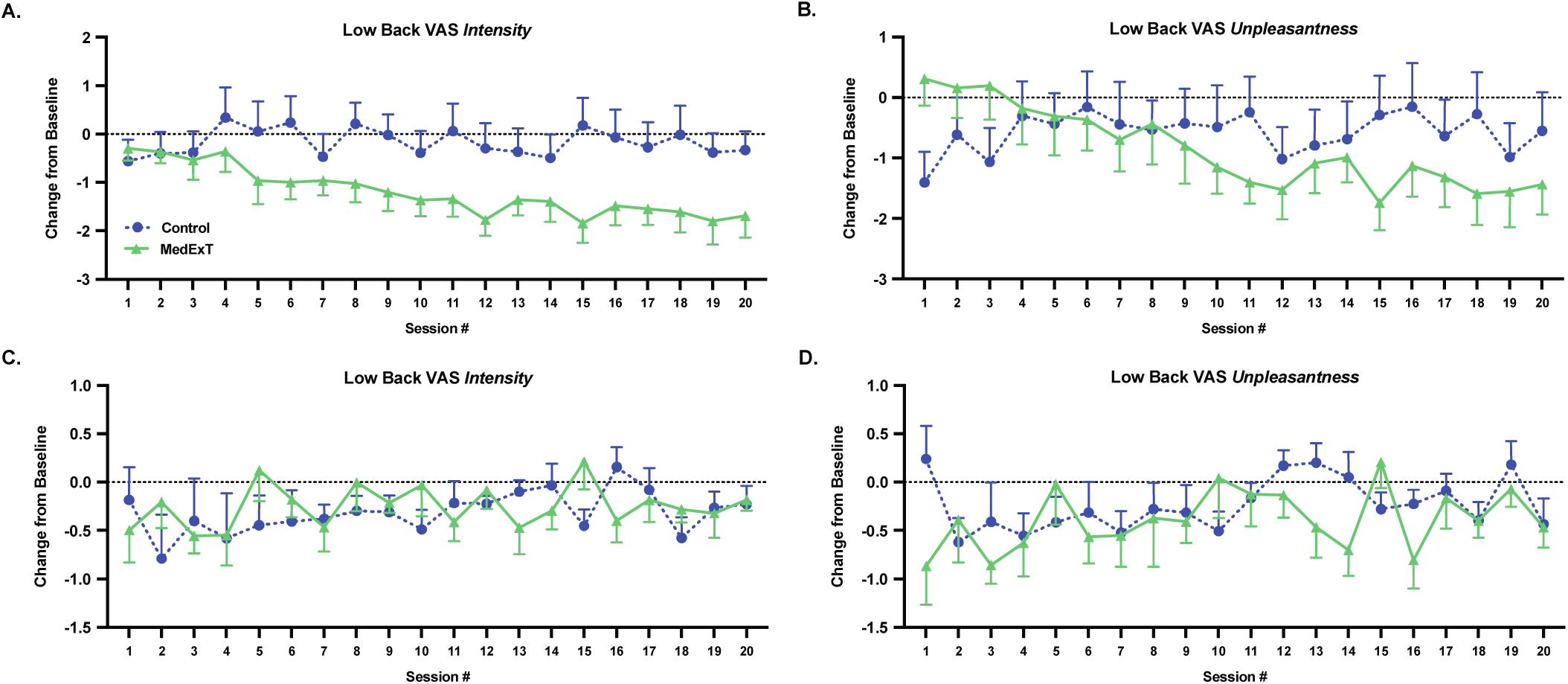
Intervention and acute effects of MedExT intervention compared to control. Data shown as mean +/− SEM with “analgesic” responses being values lower than “0” on y-axes. Intervention effects are shown in A-B comparing VAS measurement taken immediately prior to each day’s session versus the baseline VAS measurement taken on intake day. Statistically significant analgesic effects were seen in the MedExT group for (A) VAS intensity (Time (***p=0.0008), time x treatment (**p=0.0012)) and (B) VAS unpleasantness (Treatment (*p=0.0330), time (**p=0.0022), time x treatment (****p<0.0001)). Acute intervention effects shown in C-D comparing VAS taken after each day’s intervention to the VAS taken immediately before the intervention. No significant differences were found for (C) VAS intensity (n.s.) while (D) a small effect of Treatment (**p=0.0049) was found for VAS unpleasantness.

An additional measure of low back pain was assessed at the exit session. Patients were asked to recall their average pain intensity and unpleasantness before the study (i.e. at baseline) and also across the last month of being in the trial (i.e. at exit session). Two-sample t-tests revealed significant mean change differences between study groups for both intensity ratings (p=0.0167; **Fig. 5A**) and unpleasantness ratings of low back pain (p=0.0144; **Fig. 5B**) with the MedExT group showing significant improvement in their subjective evaluation of the intensity and unpleasantness of their back pain.

**Figure 5.**
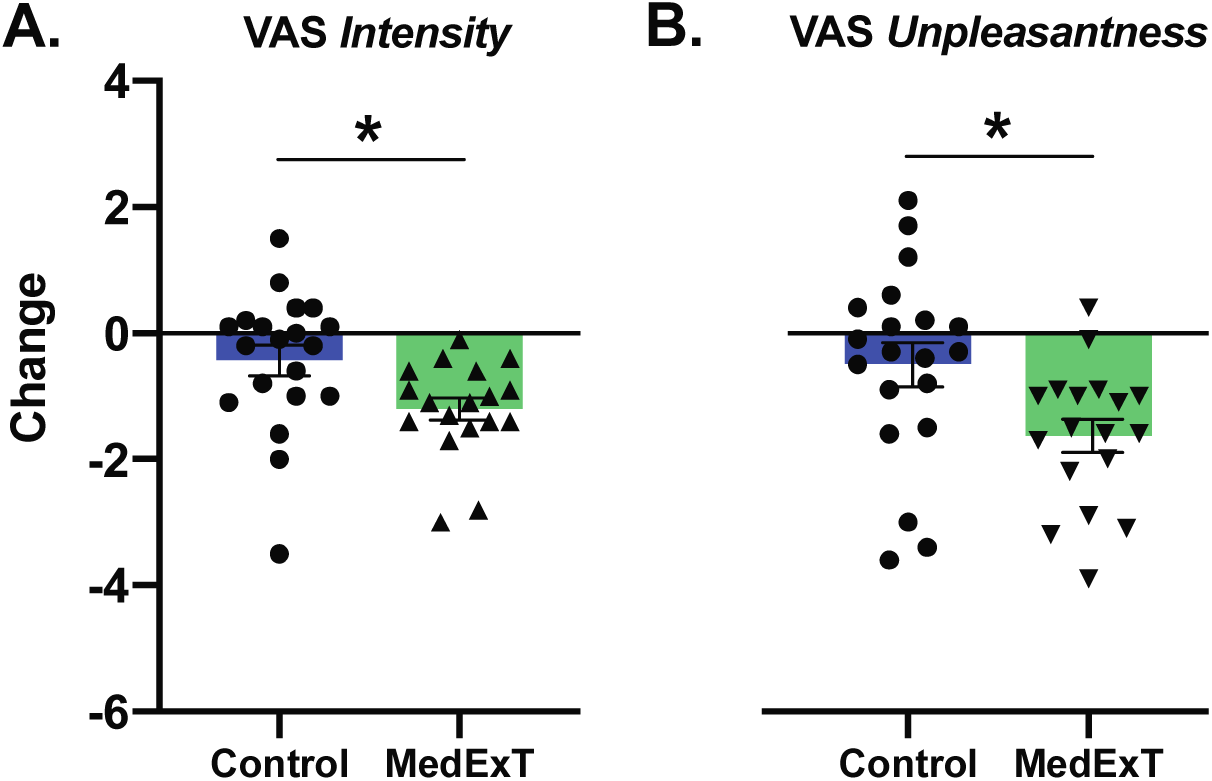
Continued patient compliance data. Shown below are the qualitative results of each outcome. (A) Box plots are shown to represent data. Box represents IQR (bottom line=Q1, middle=median, top=Q3. Whiskers represent range of data (min and max). (A) [Median=Q1 for MedExT and Control for question 1]. (B) Self-reported barriers to continued treatment are shown for the MedExT participants. (C) MedExT participants identified what the most beneficial aspect of the intervention was between three choices: Meditation only, Exercise only, or Both combined.

### Sub-analyses: Fully compliant patients only

*A priori*, we determined that we would evaluate all participants that completed 80% of each week’s sessions. This value was determined by whether participants completed the daily Qualtrics survey before/after their session. However, we reasoned that there may have been individuals in the MedExT experimental group that completed the survey, but did not actually complete the intervention. To potentially account for this non-compliance, we re-evaluated the ActiGraph GT9X watch data. We were able to monitor activity of all subjects in-clinic, as well as outside of the clinic to estimate compliance with the exercise protocol. Using walking step data from in-clinic sessions as comparison, in addition to Qualtrics survey daily log input from participants (time start and finish completed intervention), we were able to estimate participation in the walking exercise portion of the intervention for at-home sessions. We used this data to run a sub-analysis on the data. We re-ran the full data analysis on our primary and secondary outcomes for subjects that were deemed fully compliant (n=33). A list of all results is shown in **Table 3**.

**Table 3.**
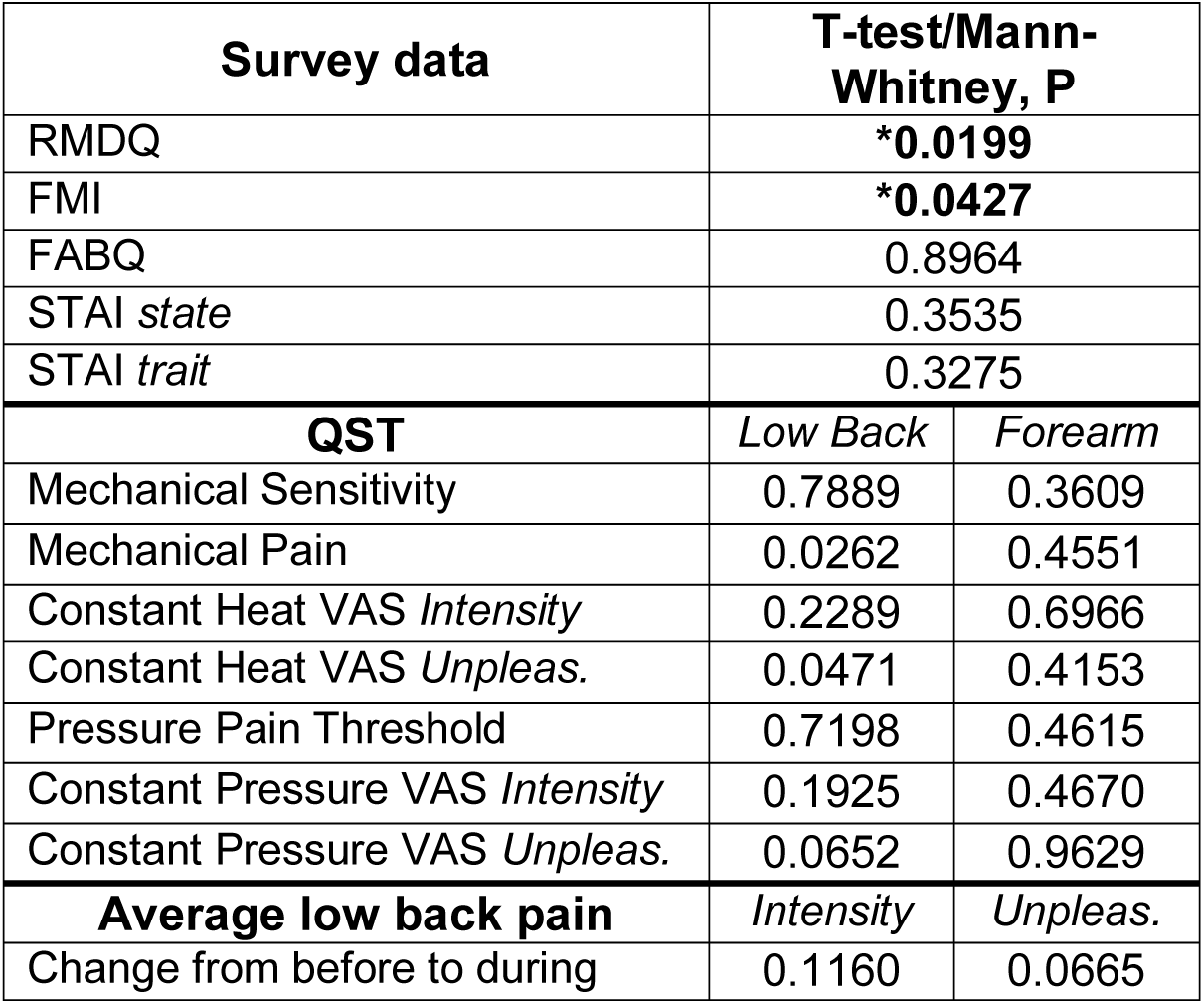
Sub-analysis of participants that completed 80% or more sessions per week.

For the primary outcome, a two-sample t-test revealed a statistically significant improvement in RMDQ scores between the MedExT and the control group (p=0.0199). FMI scores for the MedExT group significantly increased from pre to post as measured by a two-sample t-test (p=0.0427). For the psychological inventories, two-sample t-tests revealed no significant differences from pre to post for the MedExT group for FABQ (p>0.02), STAI state (p>0.02) or STAI trait (p>0.02). Similar to our full data set, no significant differences were found between groups for QST for the low back or non-dominant forearm (p>0.005). Two-sample t-test indicate no significant differences between groups for average change in low back pain ratings of intensity (p=0.1160) nor unpleasantness (p=0.0665).

### Exit Survey Data

At the exit session, all patients were asked to complete a continued patient compliance survey that we developed. The results of these outcomes are shown in **Fig. 6**. This survey sought to identify the need for pain treatments during the study (**Fig. 6A)**, the likelihood of continued compliance post-study (**Fig. 6A**), any barriers to continuing the combined treatment (**Fig. 6B**) and the most beneficial aspect of the intervention between meditation, exercise, or the combination (**Fig. 6C**). We found qualitatively that MedExT participants reported a greater decrease in pain medication use and seemed fairly likely to continue the intervention. Importantly, when MedExT experimental participants were asked to identify the most beneficial aspects of the intervention (i.e. meditation, exercise or both) a majority of participants stated that “Both” components of the intervention were the most important.

**Figure 6.**
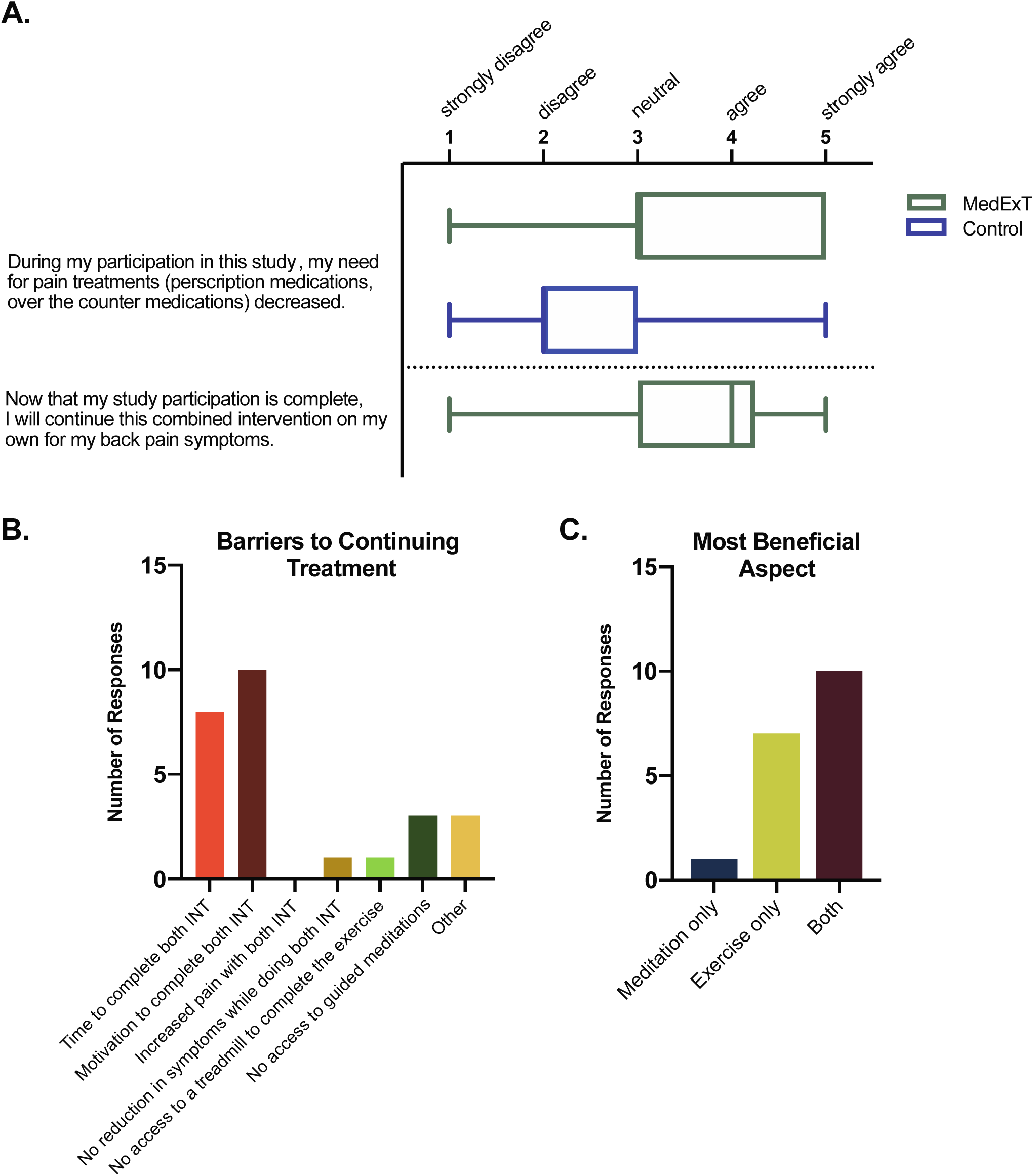
Continued patient compliance data. Shown below are the qualitative results of each outcome. (A) Box plots are shown to represent data. Box represents IQR (bottom line=Q1, middle=median, top=Q3. Whiskers represent range of data (min and max). (A) [Median=Q1 for MedExT and Control for question 1]. (B) Self-reported barriers to continued treatment are shown for the MedExT participants. (C) MedExT participants identified what the most beneficial aspect of the intervention was between three choices: Meditation only, Exercise only, or Both combined.

## DISCUSSION

In the current study, we assessed the effect of a combined intervention of mindfulness meditation followed by aerobic walking exercise in chronic low back pain. The main findings of this study indicate that meditation and exercise together were able to reduce disability, increase mindfulness and decrease self-reported ratings of low back pain. To our knowledge, this specific therapy combination has not yet been tested in chronic low back pain patients.

While the present study took a unique approach to combined mindfulness and aerobic exercise, there is a robust literature suggesting that such an approach could work. First and foremost, previous studies have tested MBSR [4, 12, 20, 38, 39, 47] and mindfulness meditation [58] alone, as well as aerobic walking exercise programs [5, 10, 18, 24, 30, 32, 40, 50] in low back pain patients. Overall, these studies found improved disability, sleep quality, psychological function, depression, affective pain perception, fitness, pain severity, and reduced need for pain medications. One important aspect of the mindfulness used in the present study was the accessibility of the mindfulness. Beyond the introductory 45-minute session with a clinical psychologist, our participants were naïve meditators. Nonetheless, using only five short recordings repeatedly, their mindfulness increased as assessed by the FMI. Gains seen in this study via the FMI compare to more intensive training exercises [11]. Although the recordings used here were curated for their emphasis on mindfulness, they were not specifically recorded for this intervention. We would hypothesize that the development of a mindfulness recording that specifically prepared participants for the subsequent exercise could be even more beneficial. The benefits seen here with a brief meditation program are consistent with more recent data showing that only 4 days of mindfulness-based mental training can reduce pain [55-57]. Importantly, these previous studies were done in healthy participants with models of acute nociception, while here we are showing gains in mindfulness in a chronic patient population.

In our previous work, we have elucidated that increasing the frequency component of exercise dose to be the most likely to have a positive effect on chronic pain patients [43]. We tested these predictions in a trial with multiple doses of moderate intensity treadmill walking exercise in healthy subjects. Treadmill walking exercise was chosen because it was easy to implement, required little to no training and had more precision in controlling dosage. In this study, we found that the moderate dose, or 5x of 30 minutes/day of this exercise regimen in one week proved to be the most optimal for reducing cutaneous pressure pain ratings [42]. We reasoned that this aerobic exercise intervention was a good starting dose for reducing pain outcomes. In addition, this prescription aligns with that of ACSM’s recommendation for physical activity for healthy individuals which is 150 MET minutes per week [21].

Surprisingly, while MedExT participants rated lower disability along with lower on-going pain and lower average pain compared to the start of trial, they failed to show any changes in fear avoidance behavior or anxiety. This result is in contrast to data generated from a similar study that implemented combined mindfulness and exercise in the context of major depression [2]. That study found 8 weeks of 60-minute twice a week mental and physical training significantly reduced depressive symptoms and ruminative thoughts. It is possible that the lack of a significant effect on fear and anxiety in the present study was driven by the lower starting anxiety and fear levels in our cohort of participants. We anticipate that the anxiolytic effects of the combined intervention may only present itself in the context of higher baseline anxiety and fear avoidance behavior or with longer duration studies (i.e. 8 vs 4 weeks).

During the trial, participants rated pain before and after each session. Thus, we were able to track their on-going pain before each intervention compared to baseline and evaluate the potential for acute analgesic effects of the intervention itself. Analysis of day to day pre-intervention ratings of on-going low back pain revealed significant effects of time and time x treatment for intensity (p=0.0144, p=0.0012) and unpleasantness (p=0.0237, p<0.0001), respectively. Acutely however, there were no significant changes comparing pre-intervention to post-intervention on single days. Taken together, these data are indicative of a time dependent effect of the intervention with beneficial outcomes resulting from cumulative repeated treatment sessions. Interestingly, the data do not begin to show separation across time between the groups until about day eight for pain intensity ratings and day ten for pain unpleasantness ratings. This suggests that for this specific intervention, sustained analgesic benefit can only be achieved after 8-10 sessions or about two weeks of effort.

Our data show significant differences in qualitative ratings of low back pain, but no significant effects with QST. Interestingly, we do see trends for decreased ratings of pain unpleasantness (p=0.0338) in response to a noxious constant pressure stimulus applied to the low back. Although, subjects’ pressure pain threshold was unchanged from baseline, their perception of that same pain declined. These paradoxical effects are consistent with that of a previous study that tested aerobic treadmill walking exercise on painful QST in healthy participants [42]. Similar effects have also been shown in sustained aerobic exercise, where training induced increases in pain tolerance, but unaltered pressure pain threshold [29].

Using daily Qualtrics survey monitoring and wrist-worn activity trackers, we were able to estimate compliance from participants beyond self-report. Results from a sub-analysis for fully compliant patients only (MedExT; n=13, Control; n=20) shows significant improvements in disability and an increase in mindfulness for MedExT subjects, which is consistent with our analysis of the full data set (MedExT; n=18, Control; n=20). Even the individuals who were estimated to be noncompliant (n=5) still exercised at least 7 days. Overall, this analysis suggests that compliance was not a major confound of the reported results.

Results of our continued compliance survey given during the exit session shows favorable outcomes for the combined meditation and exercise intervention. In particular, the finding that a majority of MedExT group participants found the most beneficial component of the intervention was the combination of the exercise and meditation suggests the potential for synergistic effects in this study. The need for pharmacological pain treatments trended to decrease for the MedExT group compared to control subjects, which suggests analgesic benefit of a solely nonpharmacological source. MedExT participants also indicated that they would be likely to continue the combined intervention on their own to manage their back-pain symptoms.

### Strengths and Limitations

Notable strengths to this study included a low risk of detection bias through blinding of the outcome assessor for QST, and a lack of confounding demographic variables. The average age of study participants was 37.6 years, which accurately represents the range of our inclusion criteria (18-60). Additionally, after subtracting intervention walking steps, there was no statistical difference in average steps per day between control and MedExT subjects. The most significant limitation to this study is the lack of all possible study arms. Without exercise-only and meditation-only groups, the hypothesis of synergy between the individual therapies cannot be specifically tested. This study had a higher risk of performance bias, due to lack of blinding of participants, which is very difficult due to the nature of the interventions. Also, the order of QST assessment was not randomized, which could contribute to additional outcome bias. These tests were performed from least invasive to most invasive, to avoid increased sensitization. Patients in this trial were mostly female (n=25) compared to male (n=13), however chronic low back pain is reported to be more prevalent among women [27]. According to our continued compliance survey, the most prominently identified barriers to continuing this treatment after conclusion of the trial included time and motivation to complete both interventions. Notably, no patients reported increased pain with both meditation and exercise. Thus, while we cannot state conclusively that there was synergy between the treatments, we can be confident that the therapies are not overtly antagonist. Other reported barriers included lack of access to a treadmill or guided meditations. Although, it is worth noting that access to additional meditations were provided to interested patients following their completion of the trial.

### Clinical Implications

The results of this study demonstrate that a combined therapy approach of mindfulness meditation followed by moderate intensity treadmill walking provides a significant benefit to disability, mindfulness and perception of low back pain. Patients in this treatment group also report less need for pain medications, and favorability for the combined approach as opposed to meditation or exercise therapy alone. This is the first study testing this treatment combination in chronic low back pain patients. Because synergistic benefits could not be definitely determined from this trial, future studies must be done to conclude the most efficacious combination of this treatment regimen. Nonetheless, we feel the potential for this combined approach to improve outcomes in chronic low back pain is high. As exercise and meditation (as practiced in this study) are low cost, easy to implement, and carry few negative side-effects, we are optimistic about the use of this or similar integrative therapy in the clinic.

## ACKNOWLEDGEMENTS

The authors would like to thank the participants who volunteered for this research. We would like to thank Allison Morgan, PAc and Kristin D’Acunto, PAc for assistance in performing back pain evaluations for eligibility screening. We would also like to thank Dana Farrel for research facilitation, Kerri Jackson for Pitt+Me coordination, Sadie Riskus for Qualtrics survey compilation and optimization and Dr. Ian Edwards for mindfulness meditation consultation. Funding for this work was supported by the National Institutes of Health through Grant Number UL1TR001857 and through a Pain Research Challenge Grant supported by the Clinical and Translational Science Institute at the University of Pittsburgh and the Virginia Kaufmann Foundation.

## REFERENCES

[1] Abraham, P., Rennert, R.C., Martin, J.R., Ciacci, J., Taylor, W., Resnick, D., Kasper, E., and Chen, C.C., The role of surgery for treatment of low back pain: insights from the randomized controlled Spine Patient Outcomes Research Trials. Surg Neurol Int, 2016. 7: p. 38.

[2] Alderman, B.L., Olson, R.L., Brush, C.J., and Shors, T.J., MAP training: combining meditation and aerobic exercise reduces depression and rumination while enhancing synchronized brain activity. Transl Psychiatry, 2016. 6: p. e726.

[3] Balady, G.J., Chaitman, B., Driscoll, D., Foster, C., Froelicher, E., Gordon, N., Pate, R., Rippe, J., and Bazzarre, T., Recommendations for cardiovascular screening, staffing, and emergency policies at health/fitness facilities. Circulation, 1998. 97(22): p. 2283–93.

[4] Banth, S. and Ardebil, M.D., Effectiveness of mindfulness meditation on pain and quality of life of patients with chronic low back pain. Int J Yoga, 2015. 8(2): p. 128–33.

[5] Barker, K.L., Dawes, H., Hansford, P., and Shamley, D., Perceived and measured levels of exertion of patients with chronic back pain exercising in a hydrotherapy pool. Arch Phys Med Rehabil, 2003. 84(9): p. 1319–23.

[6] Bender, T., Nagy, G., Barna, I., Tefner, I., Kadas, E., and Geher, P., The effect of physical therapy on beta-endorphin levels. Eur J Appl Physiol, 2007. 100(4): p. 371–82.

[7] Booth, M., Assessment of physical activity: an international perspective. Res Q Exerc Sport, 2000. 71 Suppl 2: p. 114–20.

[8] Borg, G.A., Psychophysical bases of perceived exertion. Med Sci Sports Exerc, 1982. 14(5): p. 377–81.

[9] Brach, T. 2019; Available from: https://www.tarabrach.com.

[10] Chan, C.W., Mok, N.W., and Yeung, E.W., Aerobic exercise training in addition to conventional physiotherapy for chronic low back pain: a randomized controlled trial. Arch Phys Med Rehabil, 2011. 92(10): p. 1681–5.

[11] Chandran, S., Raman, R., Kishor, M., and Nandeesh, H.P., The effectiveness of mindfulness meditation in relief of symptoms of depression and quality of life in patients with gastroesophageal reflux disease. Indian J Gastroenterol, 2019. 38(1): p. 29–38.

[12] Cherkin, D.C., Sherman, K.J., Balderson, B.H., Cook, A.J., Anderson, M.L., Hawkes, R.J., Hansen, K.E., and Turner, J.A., Effect of Mindfulness-Based Stress Reduction vs Cognitive Behavioral Therapy or Usual Care on Back Pain and Functional Limitations in Adults With Chronic Low Back Pain: A Randomized Clinical Trial. JAMA, 2016. 315(12): p. 1240–9.

[13] Chou, R., Deyo, R., Friedly, J., Skelly, A., Hashimoto, R., Weimer, M., Fu, R., Dana, T., Kraegel, P., Griffin, J., Grusing, S., and Brodt, E.D., Nonpharmacologic Therapies for Low Back Pain: A Systematic Review for an American College of Physicians Clinical Practice Guideline. Ann Intern Med, 2017. 166(7): p. 493–505.

[14] Cropley, M., Ussher, M., and Charitou, E., Acute effects of a guided relaxation routine (body scan) on tobacco withdrawal symptoms and cravings in abstinent smokers. Addiction, 2007. 102(6): p. 989–93.

[15] Desbordes, G., Negi, L.T., Pace, T.W., Wallace, B.A., Raison, C.L., and Schwartz, E.L., Effects of mindful-attention and compassion meditation training on amygdala response to emotional stimuli in an ordinary, non-meditative state. Front Hum Neurosci, 2012. 6: p. 292.

[16] Deyo, R.A., Mirza, S.K., and Martin, B.I., Back pain prevalence and visit rates: estimates from U.S. national surveys, 2002. Spine (Phila Pa 1976), 2006. 31(23): p. 2724–7.

[17] Dinler, M., Diracoglu, D., Kasikcioglu, E., Sayli, O., Akin, A., Aksoy, C., Oncel, A., and Berker, E., Effect of aerobic exercise training on oxygen uptake and kinetics in patients with fibromyalgia. Rheumatol Int, 2009. 30(2): p. 281–4.

[18] Eadie, J., van de Water, A.T., Lonsdale, C., Tully, M.A., van Mechelen, W., Boreham, C.A., Daly, L., McDonough, S.M., and Hurley, D.A., Physiotherapy for sleep disturbance in people with chronic low back pain: results of a feasibility randomized controlled trial. Arch Phys Med Rehabil, 2013. 94(11): p. 2083–92.

[19] Ehrlich, G.E., Low back pain. Bull World Health Organ, 2003. 81(9): p. 671–6.

[20] Esmer, G., Blum, J., Rulf, J., and Pier, J., Mindfulness-based stress reduction for failed back surgery syndrome: a randomized controlled trial. J Am Osteopath Assoc, 2010. 110(11): p. 646–52.

[21] Garber, C.E., Blissmer, B., Deschenes, M.R., Franklin, B.A., Lamonte, M.J., Lee, I.M., Nieman, D.C., Swain, D.P., and American College of Sports, M., American College of Sports Medicine position stand. Quantity and quality of exercise for developing and maintaining cardiorespiratory, musculoskeletal, and neuromotor fitness in apparently healthy adults: guidance for prescribing exercise. Med Sci Sports Exerc, 2011. 43(7): p. 1334–59.

[22] Gelles, D. How to Meditate. The New York Times, 2016.

[23] Geneen, L.J., Moore, R.A., Clarke, C., Martin, D., Colvin, L.A., and Smith, B.H., Physical activity and exercise for chronic pain in adults: an overview of Cochrane Reviews. Cochrane Database Syst Rev, 2017. 4: p. CD011279.

[24] Hartvigsen, J., Morso, L., Bendix, T., and Manniche, C., Supervised and non-supervised Nordic walking in the treatment of chronic low back pain: a single blind randomized clinical trial. BMC Musculoskelet Disord, 2010. 11: p. 30.

[25] Hauser, W., Klose, P., Langhorst, J., Moradi, B., Steinbach, M., Schiltenwolf, M., and Busch, A., Efficacy of different types of aerobic exercise in fibromyalgia syndrome: a systematic review and meta-analysis of randomised controlled trials. Arthritis Res Ther, 2010. 12(3): p. R79.

[26] Hilton, L., Hempel, S., Ewing, B.A., Apaydin, E., Xenakis, L., Newberry, S., Colaiaco, B., Maher, A.R., Shanman, R.M., Sorbero, M.E., and Maglione, M.A., Mindfulness Meditation for Chronic Pain: Systematic Review and Meta-analysis. Ann Behav Med, 2017. 51(2): p. 199–213.

[27] Hoy, D., Bain, C., Williams, G., March, L., Brooks, P., Blyth, F., Woolf, A., Vos, T., and Buchbinder, R., A systematic review of the global prevalence of low back pain. Arthritis Rheum, 2012. 64(6): p. 2028–37.

[28] Jacobs, T.L., Shaver, P.R., Epel, E.S., Zanesco, A.P., Aichele, S.R., Bridwell, D.A., Rosenberg, E.L., King, B.G., Maclean, K.A., Sahdra, B.K., Kemeny, M.E., Ferrer, E., Wallace, B.A., and Saron, C.D., Self-reported mindfulness and cortisol during a Shamatha meditation retreat. Health Psychol, 2013. 32(10): p. 1104–9.

[29] Jones, M.D., Booth, J., Taylor, J.L., and Barry, B.K., Aerobic training increases pain tolerance in healthy individuals. Med Sci Sports Exerc, 2014. 46(8): p. 1640–7.

[30] Kell, R.T. and Asmundson, G.J., A comparison of two forms of periodized exercise rehabilitation programs in the management of chronic nonspecific low-back pain. J Strength Cond Res, 2009. 23(2): p. 513–23.

[31] Kerr, C.E., Sacchet, M.D., Lazar, S.W., Moore, C.I., and Jones, S.R., Mindfulness starts with the body: somatosensory attention and top-down modulation of cortical alpha rhythms in mindfulness meditation. Front Hum Neurosci, 2013. 7: p. 12.

[32] Koldas Dogan, S., Sonel Tur, B., Kurtais, Y., and Atay, M.B., Comparison of three different approaches in the treatment of chronic low back pain. Clin Rheumatol, 2008. 27(7): p. 873–81.

[33] Kostek, M., Polaski, A., Kolber, B., Ramsey, A., Kranjec, A., and Szucs, K., A protocol of manual tests to measure sensation and pain in humans. J Vis Exp, 2016(118).

[34] Kroenke, K., Outcalt, S., Krebs, E., Bair, M.J., Wu, J., Chumbler, N., and Yu, Z., Association between anxiety, health-related quality of life and functional impairment in primary care patients with chronic pain. Gen Hosp Psychiatry, 2013. 35(4): p. 359–65.

[35] Lundberg, M., Grimby-Ekman, A., Verbunt, J., and Simmonds, M.J., Pain-related fear: a critical review of the related measures. Pain Res Treat, 2011. 2011: p. 494196.

[36] Majeed, M.H., Ali, A.A., and Sudak, D.M., Mindfulness-based interventions for chronic pain: Evidence and applications. Asian J Psychiatr, 2017. 32: p. 79–83.

[37] Millan, M.J., Descending control of pain. Prog Neurobiol, 2002. 66(6): p. 355–474.

[38] Morone, N.E., Greco, C.M., Moore, C.G., and et al., A mind-body program for older adults with chronic low back pain: A randomized clinical trial. JAMA Internal Medicine, 2016. 176(3): p. 329–337.

[39] Morone, N.E., Rollman, B.L., Moore, C.G., Li, Q., and Weiner, D.K., A mind-body program for older adults with chronic low back pain: results of a pilot study. Pain Med, 2009. 10(8): p. 1395–407.

[40] Murtezani, A., Hundozi, H., Orovcanec, N., Sllamniku, S., and Osmani, T., A comparison of high intensity aerobic exercise and passive modalities for the treatment of workers with chronic low back pain: a randomized, controlled trial. Eur J Phys Rehabil Med, 2011. 47(3): p. 359–66.

[41] Pedersen, L. and Fredheim, O., Opioids for chronic noncancer pain: still no evidence for superiority of sustained-release opioids. Clin Pharmacol Ther, 2015. 97(2): p. 114–5.

[42] Polaski A. M., Phelps A. L., Kostek M. C., Szucs K. A., Kolber B. K. Dose-related effects of moderate intensity walking exercise on sensitivity to pain in healthy humans in *American College of Sports Medicine*. 2019. Orlando, FL.

[43] Polaski, A.M., Phelps, A.L., Kostek, M.C., Szucs, K.A., and Kolber, B.J., Exercise-induced hypoalgesia: A meta-analysis of exercise dosing for the treatment of chronic pain. PLoS One, 2019. 14(1): p. e0210418.

[44] Pollock, M.L., Gaesser G.A., Butcher J.D., Despres, J.P., Dishman, R.K., Franklin, B.A., Garber, C.E., American College of Sports Medicine Position Stand. The recommended quantity and quality of exercise for developing and maintaining cardiorespiratory and muscular fitness, and flexibility in healthy adults. Med Sci Sports Exerc, 1998. 30(6): p. 975–91.

[45] Qualtrics. 2019: Provo, Utah, USA.

[46] Roland, M. and Morris, R., A study of the natural history of back pain. Part I: development of a reliable and sensitive measure of disability in low-back pain. Spine (Phila Pa 1976), 1983. 8(2): p. 141–4.

[47] Schmidt, S., Gmeiner, S., Schultz, C., Lower, M., Kuhn, K., Naranjo, J.R., Brenneisen, C., and Hinterberger, T., Mindfulness-based Stress Reduction (MBSR) as Treatment for Chronic Back Pain - an Observational Study with Assessment of Thalamocortical Dysrhythmia. Forsch Komplementmed, 2015. 22(5): p. 298–303.

[48] Speilberger, C.D., & Sydeman, S.J., *State-Trait Anxiety Inventory and State-Trait Anger Expression Inventory*. The Use of Psychological Testing for Treatment Planning and Outcome Assessment. 1994, Hillsdale, NJ, USA: L. Erlbaum Associates.

[49] Tang, Y.Y., Lu, Q., Fan, M., Yang, Y., and Posner, M.I., Mechanisms of white matter changes induced by meditation. Proc Natl Acad Sci U S A, 2012. 109(26): p. 10570–4.

[50] Turner, J.A., Clancy, S., McQuade, K.J., and Cardenas, D.D., Effectiveness of behavioral therapy for chronic low back pain: a component analysis. J Consult Clin Psychol, 1990. 58(5): p. 573–9.

[51] Ussher, M., Cropley, M., Playle, S., Mohidin, R., and West, R., Effect of isometric exercise and body scanning on cigarette cravings and withdrawal symptoms. Addiction, 2009. 104(7): p. 1251–7.

[52] Waddell, G., Newton, M., Henderson, I., Somerville, D., and Main, C.J., A Fear-Avoidance Beliefs Questionnaire (FABQ) and the role of fear-avoidance beliefs in chronic low back pain and disability. Pain, 1993. 52(2): p. 157–68.

[53] Walach, H., Buchheld, N., Buttenmuller, V., Kleinknecht, N., Schmidt, S., Measuring Mindfulness — The Freiburg Mindfulness Inventory (FMI). Personality and Individual Differences, 2006. 40: p. 1543–1555.

[54] White, G., Natural History and Antiquities of Selborne. 1908, Cassell: London.

[55] Zeidan, F., Adler-Neal, A.L., Wells, R.E., Stagnaro, E., May, L.M., Eisenach, J.C., McHaffie, J.G., and Coghill, R.C., Mindfulness-Meditation-Based Pain Relief Is Not Mediated by Endogenous Opioids. J Neurosci, 2016. 36(11): p. 3391–7.

[56] Zeidan, F., Emerson, N.M., Farris, S.R., Ray, J.N., Jung, Y., McHaffie, J.G., and Coghill, R.C., Mindfulness Meditation-Based Pain Relief Employs Different Neural Mechanisms Than Placebo and Sham Mindfulness Meditation-Induced Analgesia. J Neurosci, 2015. 35(46): p. 15307–25.

[57] Zeidan, F. and Vago, D.R., Mindfulness meditation-based pain relief: a mechanistic account. Ann N Y Acad Sci, 2016. 1373(1): p. 114–27.

[58] Zgierska, A.E., Burzinski, C.A., Cox, J., Kloke, J., Stegner, A., Cook, D.B., Singles, J., Mirgain, S., Coe, C.L., and Backonja, M., Mindfulness Meditation and Cognitive Behavioral Therapy Intervention Reduces Pain Severity and Sensitivity in Opioid-Treated Chronic Low Back Pain: Pilot Findings from a Randomized Controlled Trial. Pain Med, 2016. 17(10): p. 1865–1881.

